# Stimulation mapping and whole-brain modeling reveal gradients of excitability and recurrence in cortical networks

**DOI:** 10.1101/2024.02.26.581277

**Authors:** Davide Momi, Zheng Wang, Sara Parmigiani, Ezequiel Mikulan, Sorenza P. Bastiaens, Mohammad P. Oveisi, Kevin Kadak, Gianluca Gaglioti, Allison C. Waters, Sean Hill, Andrea Pigorini, Corey J. Keller, John D. Griffiths

**Affiliations:** Krembil Centre for Neuroinformatics, Centre for Addiction and Mental Health (CAMH), Toronto; Department of Psychiatry and Behavioral Sciences, Stanford University Medical Center, Stanford, California; Veterans Affairs Palo Alto Healthcare System, and the Sierra Pacific Mental Illness, Research, Education, and Clinical Center, Palo Alto, California; Wu Tsai Neuroscience Institute, Stanford, California; Department of Health sciences, Università degli studi di Milano; Institute of Medical Science, University of Toronto; Institute of Biomedical Engineering, University of Toronto; Dipartimento di Scienze Biomediche e Cliniche “L.Sacco”, Università degli Studi di Milano, Milano, Italy; Nash Family Center for Advanced Circuit Therapeutics, Icahn School of Medicine at Mount Sinai, New York, NY, USA; Department of Psychiatry, University of Toronto; Institute of Medical Sciences, University of Toronto; Dipartimento di Scienze Biomediche e Cliniche “L.Sacco”, Università degli Studi di Milano, Milano, Italy Department of biomedical, surgical and dental sciences, Università degli Studi di Milano

**Keywords:** Neuroimaging, computational neuroscience, brain stimulation, neuromodulation, connectomics, cognitive neuroscience, brain and therapeutics

## Abstract

The human brain exhibits a modular and hierarchical structure, spanning low-order sensorimotor to high-order cognitive/affective systems. What is the causal significance of this organization for brain dynamics and information processing properties? We investigated this question using rare simultaneous multimodal electrophysiology (stereotactic and scalp EEG) recordings in patients during presurgical intracerebral electrical stimulation (iES). Our analyses revealed an anatomical gradient of excitability across the cortex, with stronger iES-evoked EEG responses in high-order compared to low-order regions. Mathematical modeling further showed that this variation in excitability levels results from a differential dependence of recurrent feedback from non-stimulated regions across the anatomical hierarchy, and could be extinguished by suppressing those connections in-silico. High-order brain regions/networks thus show a more functionally integrated processing style than low-order ones, which manifests as a spatial gradient of excitability that is emergent from, and causally dependent on, the underlying hierarchical network structure.

## INTRODUCTION

The human brain constitutes a highly intricate network of interconnected regions that maintain ongoing communication, even during periods of rest^1^. Research employing functional MRI (fMRI) has shown how distant brain regions exhibit synchronized fluctuations (functional connectivity) in their spontaneous activity, giving rise to distinct spatial patterns of temporal covariances known as resting-state networks (RSNs)^2–4^. The topographic organization of the seven canonical RSNs (visual, somatomotor, dorsal attention, anterior salience, limbic, fronto-parietal, default-mode networks^5^) has now been extensively replicated and validated across multiple species and data modalities^6–8^.

Given the significance of RSNs across cognitive and clinical domains, a question of central importance for contemporary neuroscience research is how these structures emerge from their anatomical and physiological underpinnings. Significant progress has been made on the anatomical underpinnings following the discovery that the seven canonical RSNs adhere a distinctive spatial layout^9^ on both cortical and subcortical structures^10,11^. This layout encodes a hierarchical distinction^12^ between low-order networks (visual, somatomotor) associated with fundamental sensory/motor functions, and high(er)-order networks (limbic, fronto-parietal, dorsal attention, ventral attention, default-mode) associated with introspection, self-referential contemplation, and intricate cognitive processes^9,13^. However, major gaps still remain in this rich picture of functional brain organization, especially in two key areas: i) the dynamics and neurophysiology of RSN activity, and ii) how the spatiotemporally structured neural activity we conceptualize as co-ordinated RSN behavior emerges from interactions between underlying micro-/meso-scale circuit mechanisms and the macro-scale network structure of the anatomical connectome. One set of open questions that spans both of these areas concerns the variation across RSNs in their input response characteristics. It is widely believed, due mainly to analogies with task-activation studies, that each RSN plays a key role in one or more distinct neurocognitive processes^14,15^. This is perhaps most clearly evident in their conventionally assigned names (dorsal attention, visual, somatomotor, etc.). Implicit in this is also the idea that there are differences between RSNs in their information processing activities. In other words, the fact that all brain regions have widespread long-range connections not restricted to adjacent regions within the same or neighboring hierarchical levels^16,17^ suggests that the RSNs should differ systematically in how they respond to their inputs.

A compelling *modus operandi* to casually study principles of brain organization is the perturbational approach, which couples precisely targeted neurostimulation with concurrent fast (electrophysiological) neural activity recordings^18^. Recent studies employing this approach with concurrent transcranial magnetic stimulation and electroencephalography (TMS-EEG) have reported that stimulation-evoked responses exhibit a distinctive pattern of activity propagation, predominantly spreading to distal regions that are both structurally and functionally connected to the target site^19–21^. Importantly, these studies also demonstrate that stimulus-evoked activity preferentially propagates to, and exhibits sustained activity within, distal parts of the same (distributed, discontiguous) RSN that was used for the initial TMS targeting. In related work using intracerebral electrical stimulation (iES) in patients undergoing brain surgery, Veit and colleagues observed faster activation and spreading to regions within the stimulated RSN than those within non-stimulated RSNs^22^

In this study, we aim to explore how the interactions within and between different RSNs contribute to their modular and hierarchical organization. Are there qualitative differences between low-order and high-order RSNs in terms of their response to external stimulation? What is the level of cross-talk across RSNs in their stimulation responses? How necessary are these network-network interactions in determining a local brain response?

To address these questions, we analyzed brain activity patterns using simultaneous recordings of stereotactic electroencephalography (sEEG) and scalp high-density electroencephalographic (hd-EEG) data from patients undergoing pre-surgical iES. We then employed a whole-brain, connectome-based neurophysiological model for causally investigating the level of recurrence in cortical networks. Although standard analysis of noninvasive neuroimaging data can provide insight into neural processes in the human brain^23^, mathematical modeling^24^ can delve deeper into the underlying mechanisms of an observed empirical phenomena providing insight into mechanisms that are challenging to measure in vivo in humans. This study used our recent approach^25^ and combined novel analytical techniques and subject-specific mathematical models of brain stimulation. Using combined iES and simultaneous recordings of sEEG and scalp hd-EEG, we mapped the response properties of seven canonical RSNs across 323 stimulation sessions from 36 patients. By fitting connectome-based neurophysiological models to each patient’s hd-EEG data, we replicated the observed response patterns accurately. Additionally, we performed spatially specific ‘virtual dissections’ on the models, isolating the stimulated network from surrounding activity while preserving its ability to propagate and receive information internally. Our main question was whether low-order and high-order RSNs exhibit different information processing characteristics, which we take excitability levels and stimulated input response characteristics to be a reasonable (albeit coarse) proxy of. We hypothesized that activity patterns in the high-order networks would show a more informationally-integrated level of organization^26^, where feedback connections are necessary to generate the observed iES responses. Conversely, low-order networks would show more informationally-segregated behavior in their evoked activity dynamics^27^, with iES stimulation responses that are primarily dependent on intrinsic within-network activity - and thus relatively unchanged following virtual dissections. Understanding the role of recurrent feedback in shaping RSN information flow has implications for diagnostic and therapeutic strategies in psychiatry and neurology.

## RESULTS

### A gradient of excitability from low-order to high-order brain networks

We evaluated the magnitude of stimulation-evoked global brain activation in concurrent hd-EEG and sEEG recordings as an index of neuronal excitability across the seven canonical RSNs (Visual network: VN, Somatomotor network: SMN, Dorsal attention network: DAN, Anterior salience network: SN, Limbic network: LN, Fronto-parietal network: FPN, Default-mode network: DMN). The assessment of which RSN each sEEG electrode stimulation site fell within demonstrated a high spatial resolution as indicated by an average distance of 0.52 cm ± 0.22 cm to the nearest parcel centroid (Fig. 2A). When examining the hd-EEG global mean field power (GMFP), we observed three response clusters at ∼40 ms, ∼80 ms, and ∼370 ms (Fig. 2B). These peak response timings are consistent with results previously reported from invasive human electrophysiology recordings^22^. We observed a significant interaction between response timing and stimulated network (Fig 2C, top row; F(12, 927) = 2.539, p = 0.00266), indicating that the effect of stimulation on the overall response varied depending on both the timing that the response was recorded, and which network was stimulated. This significant interaction was supported by significant main effects of both response timing (F(2, 927) = 93.792, p < 2e-16) and stimulated network F(6, 927) = 3.641, p = 0.00141). Permuted Wilcoxon-Mann-Whitney U pairwise comparisons showed significant differences in AUC for: SN-SMN: W=29127.5, p<0.0001; SMN-DAN: W=8083, p=019; SMN-FPN: W=10960, p<0.0001; SMN-DMN: W=30315, p<0.0001; SMN-LN: W=15772, p=0.004; VN-FPN: W=3463.5, p=0.04; VN-DMN: W=8786, p=0.025; DMN-LN: W=13466, p=0.026.

**Fig. 1.**
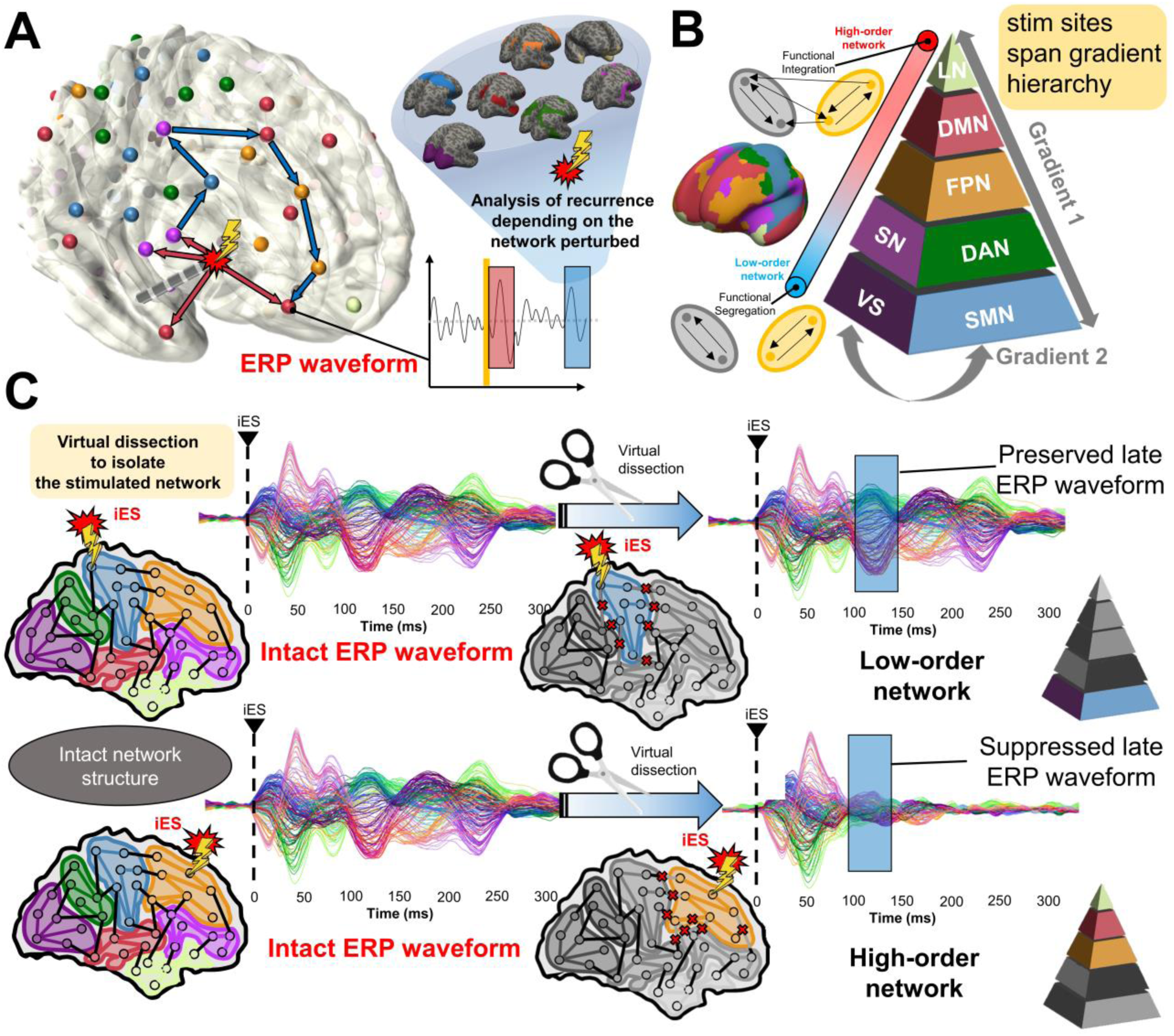
Studying RSN input processing strategies and the role of recurrent feedback with computational brain network models. Shown here is a schematic overview of the hypotheses, methodology, and general conceptual framework of the present work. **(A)** Intracerebral electrical stimulation (iES) applied to an intracortical target region generates an early (∼20-30 ms) response (evoked potential waveform component) at high-density scalp electroencephalography (hd-EEG) channels sensitive to that region and its immediate neighbors (red arrows). This also appears in more distal connected regions after a short delay due to axonal conduction and polysynaptic transmission. Subsequent second (∼60-80 ms) and third (∼140-200 ms) late evoked components are frequently observed (blue arrows). After identifying the stimulated network in this way, we aim to determine the extent to which this second component relies on intrinsic network activity versus recurrent whole-brain feedback. **(B)** Schematic of the hierarchical spatial layout of canonical resting-state networks (RSNs) as demonstrated in Margulies and colleagues^9^, spanning low-order networks showing greater functional segregation to high-order networks showing greater functional integration^12^. **(C)** Schematic of virtual dissection methodology and key hypotheses tested. We first fit personalized connectome-based computational models of iES-evoked responses to the hd-EEG time series, for each patient and stimulation location. Then, precisely timed communication interruptions (virtual dissections) were introduced to the fitted models, and the resulting changes in the iES-evoked propagation pattern were evaluated. We hypothesized that lesioning would lead to activity suppression (panel C, right side) in high-order but not low-order networks.

**Fig. 2.**
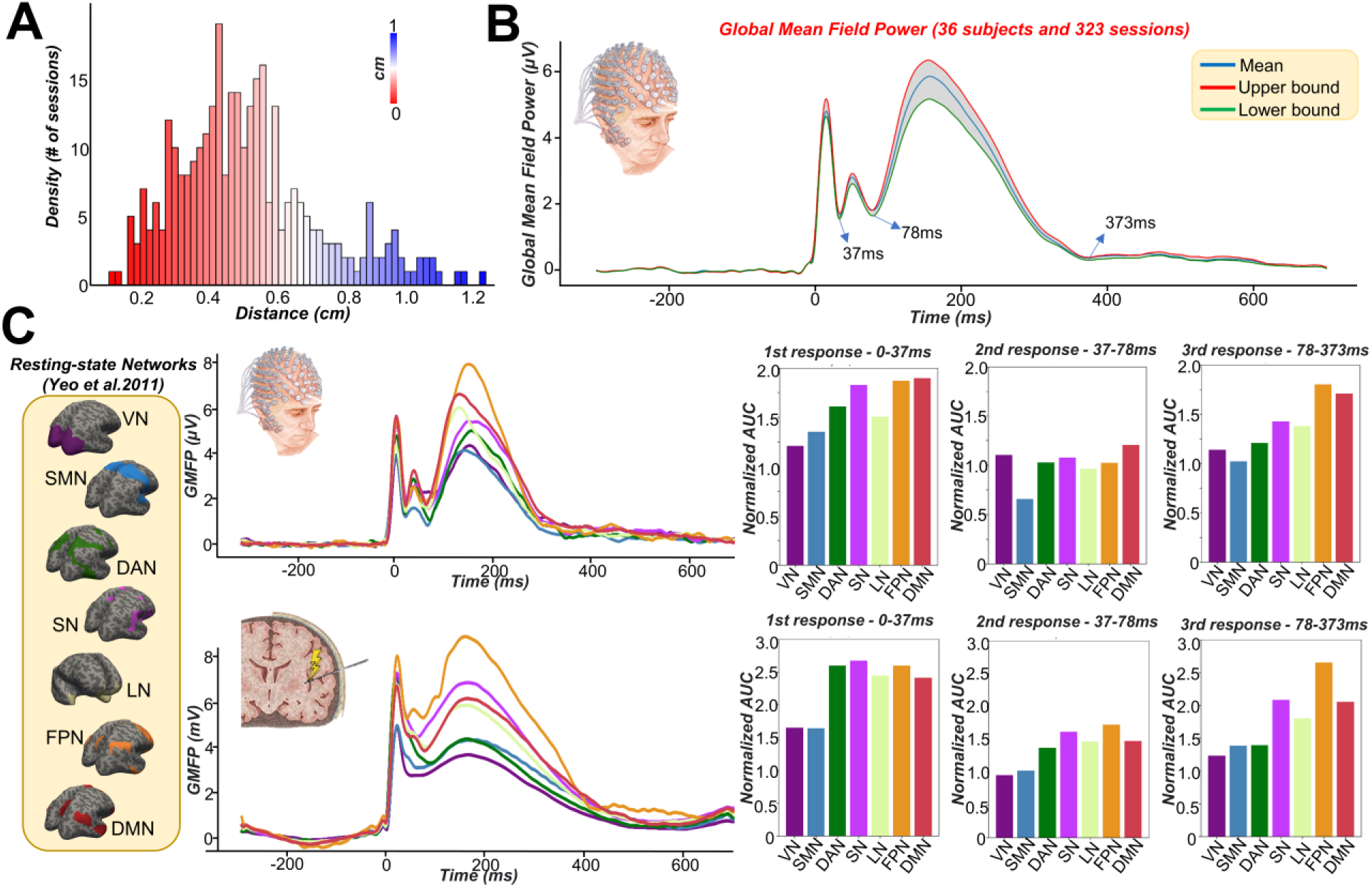
Empirical hd-EEG and sEEG signals show larger global activation patterns for high-order than low-order brain networks. **(A)** The histogram illustrates the distance in centimeters between the electrode’s centroid delivering the electrical stimulus and the center of the nearest Schaefer’s parcel^28^. The results indicate a high level of spatial precision, with the 99.9% of sessions showing distances of less than 1 cm. **(B)** Global Mean Field Power (GMFP) of hd-EEG averaged across all subjects and sessions, revealing three consistent response peaks/clusters within strict confidence intervals at ∼40 ms, ∼80 ms, and ∼370 ms, consistent with prior electrophysiological research^22^. **(C)** GMFP of every stimulated network for hd-EEG (top row) and sEEG (bottom row). Our analysis, focusing on the Area under the curve (AUC) of the three clusters, revealed a significantly stronger global activation pattern when the stimulus targeted high-order networks, such as the Default and fronto-parietal networks, particularly for the late evoked responses (third cluster at ∼370 ms). Notably, our findings mirror the ‘principal gradient’ hierarchy reported in the fMRI literature^9^, i.e.a continuous spectrum from low-order to high-order regions^12^.

When examining the sEEG data (Fig. 2C - bottom row), a significant interaction was also observed between response timing and stimulated network (F(12, 927) = 1.904, p = 0.03048). In line with the hd-EEG results, this interaction indicates that the impact of stimulation on the overall response varies depending on the network affiliation of the stimulation site. This significant interaction was supported by significant main effects in the sEEG data of both response timing (F(2, 927) = 41.961, p < 2e-16) and stimulated network F(6, 927) = 3.556, p = 0.00173). Permuted Wilcoxon-Mann-Whitney U pairwise comparisons similarly showed significant differences in AUC for: SN-SMN: W=26831, p=0.029; SN-FPN: W=9150.5, p=0.017; SMN-DAN: W=8176.5, p=0.012; SMN-FPN: W=11324, p<0.0001; SMN-DMN: W=26861, p=0.005; SMN-LN: W=16447.5, p<0.0001; VN-FPN: W=3364, p=0.014.

Overall, these findings demonstrate greater levels of excitability (as indicated by the magnitude of global activation) amongst high-order networks such as DMN and FPN, than in low-order networks such as VN or SMN. Moreover, as is also observed in both sEEG and hd-EEG data, this effect follows a continuous hierarchy over networks (Fig. 2C, right panels), that aligns closely with the well-known macroscale functional connectivity gradient ^9,13^.

### The contribution of recurrent feedback to stimulation responses mirrors the excitability gradient

Comparing the simulation runs with the intact vs. those with the lesioned structural connectome (for further details see methods), significant interactions between “response timing” (3 levels: first, second and third cluster) and “stimulated network” (F(12, 1854) = 2.397 p = 0.004490) and between “condition” (2 levels: intact, lesion) and “response timing” (F(2, 1854) = 8.798 p = 0.000157) were found.

These interactions were supported by significant main effects of “response timing” (F(2, 1854) = 268.921, p < 2e-16), “stimulated network” F(6, 1854) = 2.617, p = 0.015750) and “condition” F(1, 1854) = 11.751, p = 0.000621). Permuted Wilcoxon-Mann-Whitney U pairwise comparisons between intact and lesioned structural connectome simulations showed significant differences in AUC within the late (78-373 ms) response period for LN (W = 522.5, p = 0.01762), FPN (W = 922, p=0.00262), and DMN (W = 1352.5, p=2.994e-05). No significant differences in these late-response AUCs were found for DAN, SN, SMN and VN, and no significant differences were found for any network in the two earlier response windows (peak #1 at 0-37 ms, peak #2 at 37-78 ms). Virtual dissections applied to isolate the stimulated network thus had a significant effect on high-order networks only, and only in the later component of their stimulation responses. Moreover, as shown in Fig. 3C, while not significant for the first few, the impact of removing recurrent connections on stimulation response amplitudes follows a continuous trend from low-order to high-order networks that mirrors (in the reverse direction) the excitability gradient observed in Fig. 2.

**Fig. 3.**
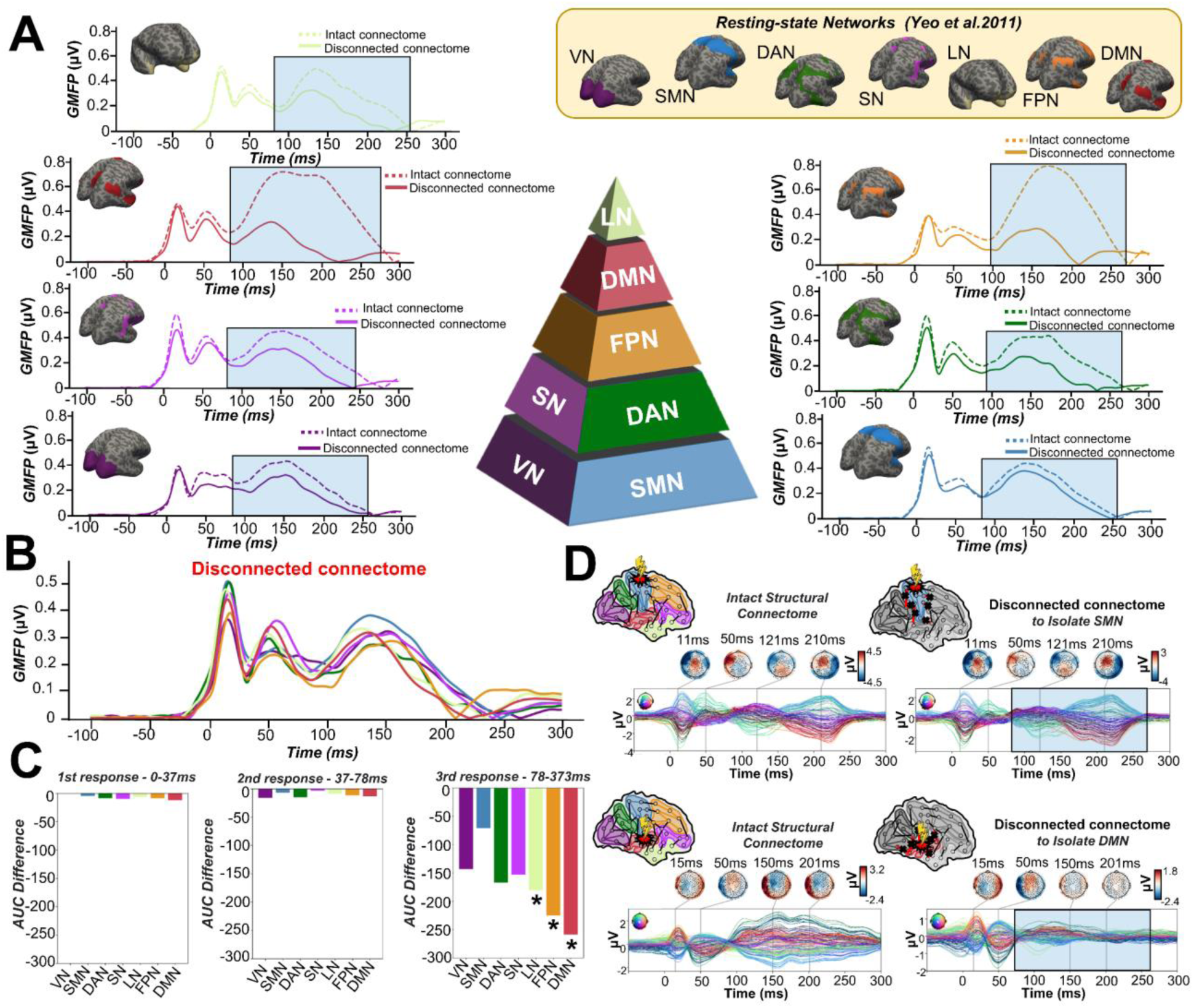
Removing recurrent connections to isolate the stimulated network suppresses late evoked-potentials for high-order networks. **(A)** Global Mean Field Power (GMFP) for every stimulated network for model-generated hd-EEG data run with both the intact (continuous line) and disconnected (dashed line) structural connectome. Findings show a more pronounced decrease in evoked late responses for high-order networks (e.g. DMN and FPN). **(B)** GMFP for every stimulated network for model-generated hd-EEG data run with the disconnected structural connectome. Unlike both empirical data and non-lesioned simulations, suppression of recurrent feedback within the model resulted in a decrease of the amplitude of the late evoked responses particularly for high-order networks (e.g. DMN, FPN). **(C)** AUC differences comparing the simulation run with the intact vs the lesioned structural connectome. A significant reduction in the AUC was found for late responses of LN, FPN and DMN compared to SMN. **(D)** Demonstration of the network recurrence-based theory for two representative sessions. Simulations of evoked dynamics are run using the intact (left) and lesioned (right) anatomical connectome. In the latter case, the connections were removed to isolate the stimulated networks for SMN (top) and DMN (bottom). In the case of the low-order network, this virtual dissection does not significantly impact the evoked potentials, while for the high-order network, a substantial reduction or disappearance of evoked components was observed. These findings indicate that, for high-order networks, the propagation dynamics depend on whole-brain integration, while for low-order networks they are mainly driven by intrinsic network reverberation.

## DISCUSSION

Using a computational framework recently developed for personalized neurostimulation modeling^25^, in this work we examined scalp and intracerebral electrophysiology data as a window into feedforward and feedback response characteristics of intrinsic human brain networks. Uncovering the rules and structures according to which brain networks are organized and interact at a mechanistic physiological level is important not only as a basic question in systems and cognitive neuroscience, but also as a foundation for clinical applications aimed at customizing brain stimulation techniques to enhance network engagement, thereby promoting better clinical outcomes. Our results demonstrate that iES leads to downstream electrophysiological evoked responses whose spatiotemporal patterning follows the hierarchical cortical gradient structure commonly studied in structural and functional neuroimaging data^9,12^. Specifically, we found significantly stronger activation patterns when the stimulus part of a high-order network (e.g. Default-mode and Fronto-parietal networks), particularly for the late evoked responses. Importantly, this trend in excitability levels was observed both in the scalp-recorded hd-EEG and the intracerebrally-recorded sEEG data, suggesting its replicability across different measurement modalities and scales of spatial resolution. Previous work has demonstrated that brain regions exhibit hierarchical gradients of activity timescales during task performance and resting state, with slower timescales found in regions most distant from sensory input and motor output^29^. These hierarchical timescales, it has been argued, serve as an intrinsic organizing principle of brain function, influencing large-scale networks and subcortical regions, across sensory and higher-order cortical regions, as well as subcortical structures.

There is growing awareness amongst neuroscientists that this hierarchical network structure of brain organization shapes the spatiotemporal propagation of activity evoked by brain stimulation^19,20,30^, and specifically that iES effects depend on the network connectivity profile of the region being stimulated^31–35^. A seminal recent study reported that patients’ self-reported perception of iES stimulation intensity depend on the stimulated region’s position in the cortical hierarchy, with simpler effects in lower-level networks and more complex, heterogeneous effects in higher-order networks^36^. Within this broader body of work, our empirical results reported here provide the first electrophysiological evidence that global patterns of hierarchical organization in the brain (cortical gradients) shape evoked-response dynamics, and specifically that the position of the stimulated region along the cortical gradient is a potent predictor iES-evoked activation dynamics.

Building on these novel observations of an excitability gradient from our empirical sEEG+hd-EEG data analyses, we used connectome-based whole-brain modeling^25^ to obtain further insights into the role of recurrent feedback activity in stimulation-evoked brain responses. Specifically, we employed a ‘virtual dissection’ approach^37^ to isolate and prevent the stimulated network from receiving feedback input from the rest of the other non-stimulated RSNs. This procedure allows us to evaluate the extent to which model-generated stimulation-evoked patterns relied on recurrent inputs from downstream brain areas that did not belong to the stimulated network. These in-silico interventions resulted in substantial reductions in the stimulation-evoked activity, with the magnitude of these reductions varying considerably depending on which network was perturbed. Virtual dissections designed to isolate the stimulated network significantly reduced the amplitude of late responses when the iES was delivered to high-order areas, as compared to low-order areas. Interestingly, in a recent work using the same virtual dissection methodology^25^, we have demonstrated that early stimulus-evoked responses are primarily driven by localized dynamics of the stimulated region, whilst later components are driven by large-scale recurrent feedback loops. In the present study, we have expanded upon these earlier findings by studying network-level responses spread widely across the cortex (as opposed to primary motor cortex stimulation only), demonstrating that these network-driven late responses differ based on the position of the stimulated region along a canonical cortical gradient hierarchy. Findings showed that the late responses mainly depend on intrinsic within-network connections for low-order regions, and extrinsic between-network connections for high-order regions. This result suggests that varying strategies are employed by different brain networks in terms of how they send, receive, and process, and is in line with other results placing sensorimotor areas (with predominantly bottom-up outgoing connections) at the bottom of the hierarchy, and higher-order association areas (with mostly top-down outgoing connections) at the top of the hierarchy^38,39^. Recent studies have also analyzed network-based incoming and outgoing communication efficiencies, characterizing low-order cortical regions as primarily senders, and high-order networks as receivers^40^. This picture is consistent with reports of a developmentally-driven shift in macroscale cortical organization during adolescence, progressing from a functional motif dominated by low-order regions (e.g. Sensorimotor, Visual) in children to an adult-like gradient, where the high-order regions are located at the opposite end of a spectrum^41^. Our findings expand this evidence base, demonstrating the existence of the macroscale functional gradients for stimulus-evoked electrophysiological data, and provide computational evidence of how this scaffold shapes information processing strategies characterized by functional segregation/integration for low-order/high-order networks respectively.

Our results, and the framework for investigating the scientific questions we are introducing here, has clear and practical relevance to basic and clinical research, as well as broader implications for the scientific understanding of functional brain organization. Using computational modeling and the virtual dissection approach allows us to ask and answer causal questions around the necessity and sufficiency of various anatomical and physiological components in different aspects of local and global brain dynamics. It also provides a potential entry point for understanding brain disorders at a causal mechanistic level, possibly leading to novel, more effective therapeutic interventions.

## MATERIALS AND METHODS

The analyses conducted in the present study consist of four main components: (i) measurement of stimulation-evoked responses in sEEG and hd-EEG data, (ii) construction of anatomical connectivity priors for our computational model using diffusion-weighted MRI tractography, (iii) simulation of whole-brain dynamics and stimulation-evoked responses with a connectome-based neural mass model, and (iv) fitting of the model to individual-subject scalp hd-EEG data. A schematic overview of the overall approach is given in Fig. 4.

**Fig. 4.**
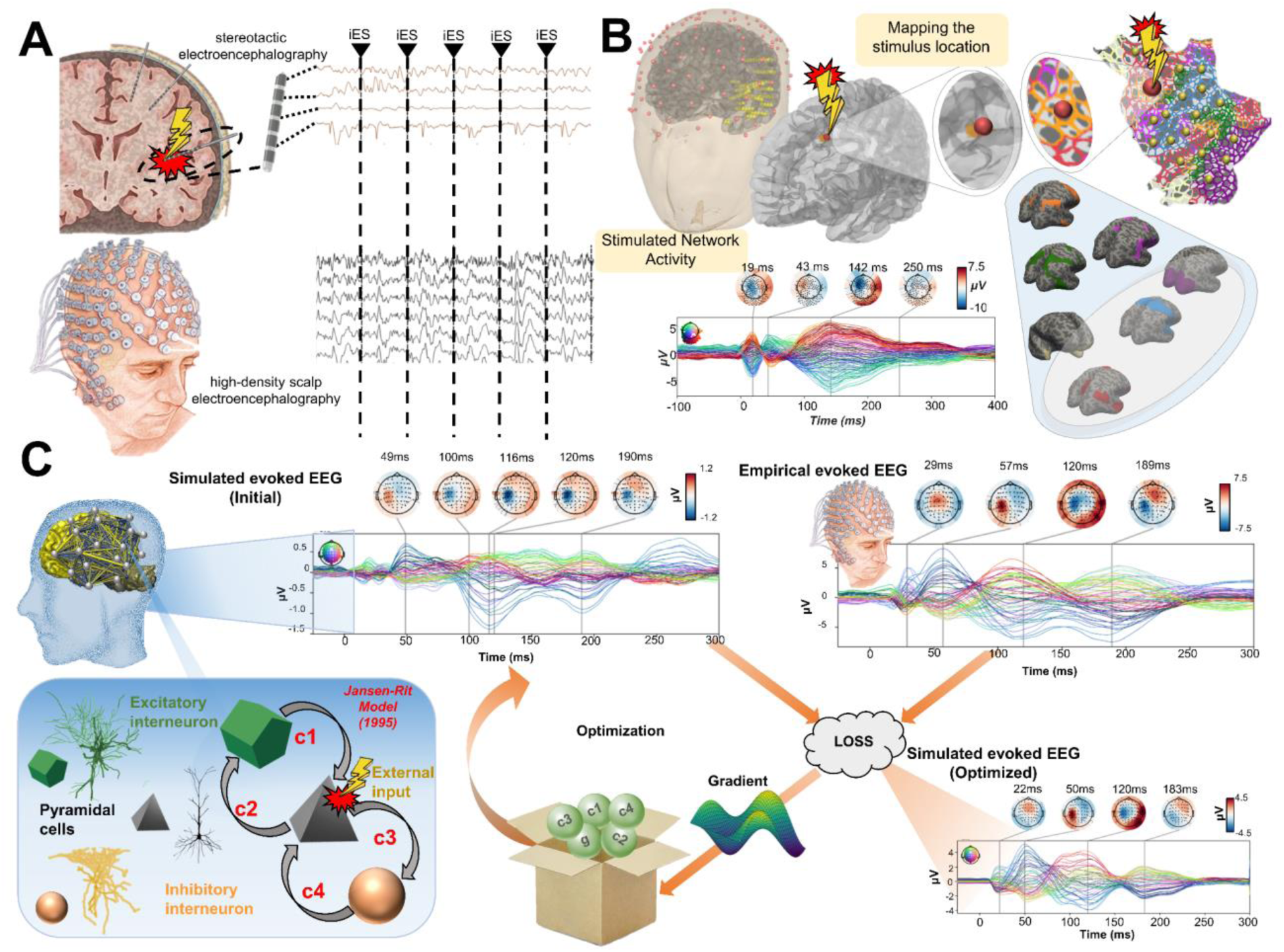
Methodological workflow for characterizing the stimulated network and performing subject-specific connectome-based neurophysiological modeling of evoked potentials. **(A)** Simultaneous stereotactic electroencephalography (sEEG) and scalp high-density electroencephalography (hd-EEG) signals were recorded. The black triangle and dashed vertical line indicate the time at which iES was delivered. For further details on the methodology and data preprocessing please refer to ^42,43^ **(B)** To pinpoint the brain network where the stimulus was delivered, we employed the Schaefer atlas^28^, which divides the brain into 1000 regions across seven distinct RSNs: visual, somatosensory, limbic, dorsal attention, ventral attention, fronto-parietal and default mode. Subsequently, we identified the parcellation region that overlapped with the intracerebral electrode responsible for delivering the stimulus, ultimately enabling us to determine the stimulated network. **(C)** To model-individual stimulus-evoked time series, the Jansen-Rit model^44^, a neural mass model comprising pyramidal, excitatory interneuron, and inhibitory interneuron populations, was embedded in every parcel of the lower-resolution 200-region Schaefer atlas^28^ for simulating and fitting neural activity time series. The connectivity between regions was modeled using diffusion-weighted MRI tractography computed from a sample of healthy young individuals from the Human Connectome Project (HCP) Dataset^45^, and then averaged to give a grand-mean anatomical connectome. The iES-induced depolarization of the resting membrane potential was modeled by a perturbing voltage offset to the mean membrane potential of the excitatory interneuron population. Next, a lead field matrix was employed to project the time series from the cortical surface parcels into EEG channel space, resulting in the generation of simulated scalp hd-EEG measurements. The quality of fit (loss) was quantified by calculating the cosine similarity between the simulated and empirical stimulus-evoked time series. Optimization of model parameters was accomplished by leveraging the autodiff-computed gradient^46^ between the objective function and the model parameters, employing the ADAM algorithm^47^. Ultimately, the optimized model parameters were utilized to generate the fitted, simulated (optimized) stimulus-evoked hd-EEG activity.

### Simultaneous stereo and high-density EEG data

The data used in this study were taken from an open dataset collected at the Claudio Munari Epilepsy Surgery Center, Milan (https://doi.org/10.17605/OSF.IO/WSGZP), where sEEG and scalp hd-EEG was recorded following single-pulse intracerebral electrical stimulation (iES) on 36 patients (median age = 33 ± 8 years, 21 female). All subjects had a history of drug-resistant, focal epilepsy, and were candidates for surgical removal/ablation of the seizure onset zone (SOZ). For details regarding the data acquisition and the preprocessing steps please refer to the original papers^42,43^. All the preprocessed sEEG and hd-EEG analyses were performed using the MNE software library^48^ (mne.tools/stable/index.html) running in Python 3.6.

### Precise identification of the stimulated network

In order to identify the network stimulated for a specific session (Fig. 4B), The Schaefer atlas^28^, which divides the brain into seven canonical functional brain networks, subdivided at multiple spatial scales (100, 200, 300…1000 parcels), was mapped to the individual’s FreeSurfer parcellation. In this study we used finest-resolution (1000 region per hemisphere) Schaefer parcellation for categorizing surgical stimulation sites, and a lower-resolution (100 regions per hemisphere) for whole-brain physiological modeling and network analysis.

The seven canonical networks correspond to: Visual network: VN, Somatomotor network: SMN, Dorsal attention network: DAN, Anterior salience network: SN, Limbic network: LN, Fronto-parietal network: FPN, Default mode network: DMN. We first projected the seven-network cortical atlas onto the subject’s cortical surface using the Freesurfer spherical registration parameters. The resulting maps were then resampled to native space structural T1w MRIs. Then, we identified the parcellation region overlapping with the intracerebral electrode responsible for delivering the stimulus, ultimately allowing us to determine the stimulated network.

### Analyzing differences in the activation dynamics dependent on the stimulated network

All statistical analyses were carried out using R version 2023.06.2, Build 561. We aimed to investigate whether the pattern of activation dynamics resulting from iES depend on the specific network that is stimulated. In order to explore this, the global mean field power (GMFP) was extracted from every stimulation session.

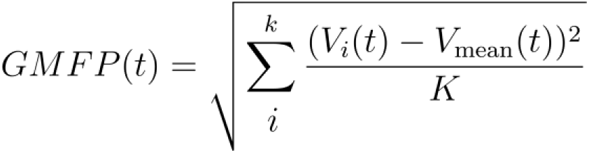

where *t* is time, V_i_(t) is the voltage at channel *i,* channel i time *t, V_mean_* is the mean of the voltage over all channels, and *K* is the number of channels. Upon examining the average scalp hd-EEG GMFP across subjects and sessions, we identified three clusters of response peaks in a time frame consistent with findings already reported in electrophysiological data from previous research using similar approaches^22^. We extracted the area under the curve (AUC) - which reflects cortical excitability^49,50^ - for each one of these clusters (Fig. 3B; cluster 1: 0 to 37 ms; cluster 2: 37 to 78 ms; cluster 3: 78 to 373 ms), and subsequently grouped the AUC values belonging to the same stimulated network session. This allowed us to assess whether the overall activity evoked by the stimulation varies systematically as a function of the specific network that was perturbed. In order to account for the varying number of sessions among participants, a mixed-design analysis of variance (ANOVA) was conducted with “response timing” as a within-subjects factor corresponding to the 3 response clusters (3 levels: first, second and third cluster) and “stimulated network’’ as a between-subjects factor corresponding to the seven networks (7 level: VN, SMN, DAN, SN, LN, FPN, DMN).

A Wilcoxon-Mann-Whitney U test was then conducted to evaluate pairwise comparisons between the different stimulated networks. Each comparison was assessed with a null distribution constructed from 1000 random permutations, with a significance threshold set at p<0.05. By comparing these conditions, we sought to determine statistically significant network-wise differences in AUC, without making any assumptions about the underlying distribution of the data.

### Overview of computational modeling approach

We employed a whole-brain modeling^51^ approach to analyze hd-EEG data and study the physiological mechanisms of network excitability. The specific model we used here incorporated 200 distinct brain regions (as defined by the Schaefer 200 parcellation), connected with a set of inter-regional weights derived from the anatomical connectome. Jansen-Rit neural mass dynamics^44^ at each region described the process of stimulated activation and oscillatory responses resulting from local interactions within cortical microcircuits, with these effects propagating to regions distal to the stimulated site via the long-range anatomical connections. After specifying its structure and a common set of priors for all parameters, the model was fit to EEG data separately for each patient. This resulted in a set of individualized physiological and connectivity parameters, having mechanistic causal influence on several spatial and temporal features of the brain stimulation response, which we subsequently interrogated to obtain further insight into our research questions around topographic organization and network specificity. For details regarding the computational model and the parameter estimation see ^25,52^ and supplementary material and methods. For a graphical overview of all optimized parameters and their distributions, see supplementary Fig. S1.

### Assessing the similarity between simulated and empirical evoked responses

To further assess the goodness-of-fit of the simulated waveforms arrived at after convergence of the ADAM algorithm, Pearson correlation coefficients and corresponding p-values between empirical and model-generated waveforms were computed for each subject. In order to control for type I error, this result was compared with a null distribution constructed from 1000 time-wise random permutations, with a significance threshold set at p<0.05.

### Dissecting the network-specific activation dynamics

The primary objective of this study is to determine the extent to which the activation patterns observed in sEEG and hd-EEG data depend on intrinsic dynamics within the stimulated network, or on contributions from other non-stimulated network regions. In order to explore this, simulations were re-run for each subject using their optimal parameters estimated from the original evoked-potentials fitting step, but this time with specifically-designed ‘virtual dissections’ applied to the (otherwise intact) structural connectome. These virtual dissections were performed by setting to zero the weights of all connections returning to the stimulated network from other non-stimulated RSNs. In this way the stimulated network was still able to send information to the whole-brain, and receive information from regions that belong to the same network. Once the whole-brain model was re-run with these new virtually dissected connectome weights, the evoked potential time series of each brain region were again projected to the hd-EEG channel space, and the AUC was extracted for the same clusters, and compared against the original model-generated evoked potentials’ AUC. For this comparison, a mixed-design ANOVA was run with “condition” as a within-subjects factor, corresponding to the 2 simulation runs with different connectomes (2 levels: intact, lesion); “response timing” as a within-subjects factor, corresponding to the 3 response clusters (3 levels: first, second and third cluster); and “stimulated network’’ as a between-subjects factor, corresponding to the seven Yeo networks (7 levels: VN, SMN, DAN, SN, LN, FPN, DMN). Then, the Wilcoxon-Mann-Whitney U test was conducted to evaluate pairwise differences between the two simulation conditions across different stimulated networks. Every comparison was compared with a null distribution constructed from 1000 time-wise random permutations, with a significance threshold set at p<0.05. We hypothesized that when the stimulus is delivered to high-order networks, these virtual dissections will significantly suppress later responses, as the activity of these networks is intricately integrated and heavily reliant on recurrent feedback from the rest of the brain. In contrast, we expect the propagation dynamics when the stimulus is delivered to low-order RSNs to remain largely unaltered, due to the fact that their activity is characterized by segregated communication strategies.

### Code availability

Full code for reproduction of the data analysis and model fitting described in this paper is freely available online at https://github.com/Davi1990/Momi_et_al_2024 and https://github.com/griffithslab/whobpyt.

### Data availability

As noted above, sEEG and hd-EEG data were taken from an open dataset publicly available at the EBRAINS platform (https://ebrains.eu/) and at Open Science Framework (https://doi.org/10.17605/OSF.IO/WSGZP). The dataset is provided in BIDS format^53^ and includes: simultaneous hd-EEG and sEEG from a total of 323 iES sessions, obtained from 36 subjects. In addition, it includes the spatial locations of the stimulating contacts in native MRI space, MNI152 space and Freesurfer’s surface space, as well as the digitized positions of the 185 scalp hd-EEG electrodes. It also contains the MRI of each subject, de-identified with AnonyMi^54^. Structural MRI data used in this study for specifying anatomical connectivity priors are available from the original Human Connectome Project dataset^45^, and have been used for similar purposes in previous work^25^.

## REFERENCES

1. Biswal, B., Yetkin, F. Z., Haughton, V. M. & Hyde, J. S. Functional connectivity in the motor cortex of resting human brain using echo-planar MRI. Magn. Reson. Med. 34, 537–541 (1995).

2. Fox, M. D. et al. The human brain is intrinsically organized into dynamic, anticorrelated functional networks. Proc. Natl. Acad. Sci. U. S. A. 102, 9673–9678 (2005).

3. Fox, M. D., Snyder, A. Z., Zacks, J. M. & Raichle, M. E. Coherent spontaneous activity accounts for trial-to-trial variability in human evoked brain responses. Nat. Neurosci. 9, 23–25 (2006).

4. Raichle, M. E. & Snyder, A. Z. A default mode of brain function: a brief history of an evolving idea. NeuroImage 37, 1083–1090; discussion 1097-1099 (2007).

5. Yeo, B. T. T. et al. The organization of the human cerebral cortex estimated by intrinsic functional connectivity. J. Neurophysiol. 106, 1125–1165 (2011).

6. Kucyi, A. et al. Intracranial Electrophysiology Reveals Reproducible Intrinsic Functional Connectivity within Human Brain Networks. J. Neurosci. 38, 4230–4242 (2018).

7. Foster, B. L., Rangarajan, V., Shirer, W. R. & Parvizi, J. Intrinsic and Task-Dependent Coupling of Neuronal Population Activity in Human Parietal Cortex. Neuron 86, 578–590 (2015).

8. Nir, Y. et al. Coupling between Neuronal Firing Rate, Gamma LFP, and BOLD fMRI Is Related to Interneuronal Correlations. Curr. Biol. 17, 1275–1285 (2007).

9. Margulies, D. S. et al. Situating the default-mode network along a principal gradient of macroscale cortical organization. Proc. Natl. Acad. Sci. 113, 12574–12579 (2016).

10. Hori, Y. et al. Cortico-Subcortical Functional Connectivity Profiles of Resting-State Networks in Marmosets and Humans. J. Neurosci. 40, 9236–9249 (2020).

11. Li, J. et al. Mapping the subcortical connectivity of the human default mode network. NeuroImage 245, 118758 (2021).

12. Mesulam, M. M. From sensation to cognition. Brain J. Neurol. 121 (Pt 6), 1013–1052 (1998).

13. Leech, R. et al. Variation in spatial dependencies across the cortical mantle discriminates the functional behaviour of primary and association cortex. Nat. Commun. 14, 5656 (2023).

14. Finn, E. S. et al. Functional connectome fingerprinting: identifying individuals using patterns of brain connectivity. Nat. Neurosci. 18, 1664–1671 (2015).

15. Greicius, M. D., Krasnow, B., Reiss, A. L. & Menon, V. Functional connectivity in the resting brain: A network analysis of the default mode hypothesis. Proc. Natl. Acad. Sci. U. S. A. 100, 253–258 (2003).

16. Honey, C. J. et al. Predicting human resting-state functional connectivity from structural connectivity. Proc. Natl. Acad. Sci. 106, 2035–2040 (2009).

17. Jung, J., Cloutman, L. L., Binney, R. J. & Lambon Ralph, M. A. The structural connectivity of higher order association cortices reflects human functional brain networks. Cortex 97, 221–239 (2017).

18. Paus, T. Mapping brain maturation and cognitive development during adolescence. Trends Cogn. Sci. 9, 60–68 (2005).

19. Momi, D. et al. Network-level macroscale structural connectivity predicts propagation of transcranial magnetic stimulation. NeuroImage 229, 117698 (2021).

20. Momi, D. et al. Perturbation of resting-state network nodes preferentially propagates to structurally rather than functionally connected regions. Sci. Rep. 11, 12458 (2021).

21. Ozdemir, R. A. et al. Individualized perturbation of the human connectome reveals reproducible biomarkers of network dynamics relevant to cognition. Proc. Natl. Acad. Sci. 117, 8115–8125 (2020).

22. Veit, M. J., et al. Temporal order of signal propagation within and across intrinsic brain networks. Proc. Natl. Acad. Sci. 118, e2105031118 (2021).

23. Buzsáki, G., Anastassiou, C. A. & Koch, C. The origin of extracellular fields and currents — EEG, ECoG, LFP and spikes. Nat. Rev. Neurosci. 13, 407–420 (2012).

24. David, O. & Friston, K. J. A neural mass model for MEG/EEG: coupling and neuronal dynamics. NeuroImage 20, 1743–1755 (2003).

25. Momi, D., Wang, Z. & Griffiths, J. D. TMS-evoked responses are driven by recurrent large-scale network dynamics. eLife 12, e83232 (2023).

26. Shine, J. M. et al. The Dynamics of Functional Brain Networks: Integrated Network States during Cognitive Task Performance. Neuron 92, 544–554 (2016).

27. Cohen, J. R. & D’Esposito, M. The Segregation and Integration of Distinct Brain Networks and Their Relationship to Cognition. J. Neurosci. Off. J. Soc. Neurosci. 36, 12083–12094 (2016).

28. Schaefer, A. et al. Local-Global Parcellation of the Human Cerebral Cortex from Intrinsic Functional Connectivity MRI. Cereb. Cortex N. Y. N 1991 28, 3095–3114 (2018).

29. Raut, R. V., Snyder, A. Z. & Raichle, M. E. Hierarchical dynamics as a macroscopic organizing principle of the human brain. Proc. Natl. Acad. Sci. 117, 20890–20897 (2020).

30. Momi, D. et al. Phase-dependent local brain states determine the impact of image-guided TMS on motor network EEG synchronization. J. Physiol. n/a, (2021).

31. Shine, J. M. et al. Distinct Patterns of Temporal and Directional Connectivity among Intrinsic Networks in the Human Brain. J. Neurosci. Off. J. Soc. Neurosci. 37, 9667–9674 (2017).

32. Keller, C. J. et al. Intrinsic functional architecture predicts electrically evoked responses in the human brain. Proc. Natl. Acad. Sci. U. S. A. 108, 10308–10313 (2011).

33. Keller, C. J. et al. Induction and Quantification of Excitability Changes in Human Cortical Networks. J. Neurosci. 38, 5384–5398 (2018).

34. Khambhati, A. N. et al. Functional control of electrophysiological network architecture using direct neurostimulation in humans. Netw. Neurosci. Camb. Mass 3, 848–877 (2019).

35. Alhourani, A. et al. Network effects of deep brain stimulation. J. Neurophysiol. 114, 2105–2117 (2015).

36. Fox, K. C. R. et al. Intrinsic network architecture predicts the effects elicited by intracranial electrical stimulation of the human brain. Nat. Hum. Behav. 4, 1039–1052 (2020).

37. Aerts, H., Fias, W., Caeyenberghs, K. & Marinazzo, D. Brain networks under attack: robustness properties and the impact of lesions. Brain 139, 3063–3083 (2016).

38. Felleman, D. J. & Van Essen, D. C. Distributed hierarchical processing in the primate cerebral cortex. Cereb. Cortex N. Y. N 1991 1, 1–47 (1991).

39. Markov, N. T. et al. Anatomy of hierarchy: feedforward and feedback pathways in macaque visual cortex. J. Comp. Neurol. 522, 225–259 (2014).

40. Seguin, C., Razi, A. & Zalesky, A. Inferring neural signalling directionality from undirected structural connectomes. Nat. Commun. 10, 4289 (2019).

41. Dong, H.-M., Margulies, D. S., Zuo, X.-N. & Holmes, A. J. Shifting gradients of macroscale cortical organization mark the transition from childhood to adolescence. Proc. Natl. Acad. Sci. 118, e2024448118 (2021).

42. Parmigiani, S. et al. Simultaneous stereo-EEG and high-density scalp EEG recordings to study the effects of intracerebral stimulation parameters. Brain Stimulat. 15, 664–675 (2022).

43. Mikulan, E. et al. Simultaneous human intracerebral stimulation and HD-EEG, ground-truth for source localization methods. Sci. Data 7, 127 (2020).

44. Jansen, B. H. & Rit, V. G. Electroencephalogram and visual evoked potential generation in a mathematical model of coupled cortical columns. Biol. Cybern. 73, 357–366 (1995).

45. Van Essen, D. C. et al. The Human Connectome Project: a data acquisition perspective. NeuroImage 62, 2222–2231 (2012).

46. Rall, I. B. & Rall, L. B. Automatic Differentiation: Techniques and Applications. (Springer-Verlag, 1981).

47. Kingma, D. P. & Ba, J. Adam: A Method for Stochastic Optimization. ArXiv14126980 Cs (2017).

48. Gramfort, A. et al. MNE software for processing MEG and EEG data. NeuroImage 86, 446–460 (2014).

49. Komssi, S., Kähkönen, S. & Ilmoniemi, R. J. The effect of stimulus intensity on brain responses evoked by transcranial magnetic stimulation. Hum. Brain Mapp. 21, 154–164 (2004).

50. Komssi, S. & Kähkönen, S. The novelty value of the combined use of electroencephalography and transcranial magnetic stimulation for neuroscience research. Brain Res. Rev. 52, 183–192 (2006).

51. Griffiths, J. D., Bastiaens, S. P. & Kaboodvand, N. Whole-Brain Modelling: Past, Present, and Future. Adv. Exp. Med. Biol. 1359, 313–355 (2022).

52. Griffiths, J. D. et al. Deep Learning-Based Parameter Estimation for Neurophysiological Models of Neuroimaging Data. 2022.05.19.492664 Preprint at 10.1101/2022.05.19.492664 (2022).

53. Appelhoff, S. et al. MNE-BIDS: Organizing electrophysiological data into the BIDS format and facilitating their analysis. J. Open Source Softw. 4, 1896 (2019).

54. Mikulan, E. et al. A comparative study between state-of-the-art MRI deidentification and AnonyMI, a new method combining re-identification risk reduction and geometrical preservation. Hum. Brain Mapp. 42, 5523–5534 (2021).

