## Supplementary for "Stimulation mapping and whole-brain modeling reveal gradients of excitability and recurrence in cortical networks"

**Running title:** Stimulation mapping and whole-brain modeling reveal hierarchical network organization
**Financial disclosures:** All authors report no conflict of interest
**Keywords:** Neuroimaging, computational neuroscience, brain stimulation, neuromodulation, connectomics, cognitive neuroscience, brain and therapeutics

**Corresponding author:**
Davide Momi
Whole Brain Modelling Group
Krembil Centre for Neuroinformatics - CAMH
250 College St., Toronto, ON M5T 1R8


*Contributed equally

**1. SUPPLEMENTARY MATERIAL AND METHODS**

1.1 Frequency-domain Analyses

1.2 Neuroimaging data and definition of connectome weight priors

1.3 Large-scale connectome-based neurophysiological brain network model

1.4 Individual-subject Jansen-Rit connectome model parameter estimation from hd-EEG data

**2. SUPPLEMENTARY RESULTS**

2.1 Network differences in evoked power are associated with late low frequency

2.2 Connectome-based neurophysiological modelling accurately reproduces subject-specific evoked dynamics

**3. SUPPLEMENTARY DISCUSSIONS**

**4. SUPPLEMENTARY REFERENCES**

### 1. SUPPLEMENTARY MATERIAL AND METHODS

#### 1.1 Frequency-domain Analyses

At each of the recording sites, we computed the stimulus-evoked spectral power across a range of frequencies (2Hz-50Hz), and assess how these responses differed as a function of which network was perturbed. For each frequency of interest, we created a Morlet wavelet and convolved it with the hd-EEG data for each channel. We then calculated the power spectrum and logarithmically transformed the results. The final step involved averaging these power values over trials within the defined analysis time window (0-300ms), chosen due to the drop-off in the evoked response after 300ms. For relative power, we normalized the values using the baseline power computed during the baseline window (-300-0ms).

Once we calculated the time-frequency power spectrum, we aimed to investigate the effect of the stimulation. For this purpose, the time-frequency power spectrum of the evoked period (0-300ms) was statistically compared with its baseline window (-300-10ms) using both a condition-wise permutation testing and a cluster-based thresholding[1] as a correction for multiple comparisons. Specifically, the permutation test transformed the difference between the evoked and the baseline windows into a z-value with respect to a null distribution of surrogate conditions difference values, obtained by swapping window labels at each of 1000 permutations. The resulting z-scores were thresholded at $p < 0.05$. After conducting an extra 1000 iterations of the permutation test, a distribution of cluster sizes representing consecutive significant points under the null hypothesis of no difference was calculated. Only the time-frequency clusters surpassing the 95th percentile of this distribution were kept. This comprehensive approach enabled us to rigorously assess the impact of stimulation on the evoked time-frequency power spectrum, identifying statistically significant changes and providing valuable insights into the dynamics of neural responses to network-specific perturbation.

##### 1.2 Neuroimaging data and definition of connectome weight priors

To establish anatomical connectivity priors representative of the population, we conducted DW-MRI tractography reconstructions on a large sample of healthy young individuals (see also [2]). This dataset comprised structural neuroimaging data from 400 healthy young subjects (170 males, aged 21–35 years), sourced from the Human Connectome Project (HCP) Dataset (available at humanconnectome.org/study/hcp-young-adult)[3].

Our DW-MRI preprocessing workflow was executed on Ubuntu 18.04 LTS using tools from the FMRIB Software Library (FSL 5.0.3; https://www.fmrib.ox.ac.uk/fsl)[4], MRtrix3 (https://www.MRtrix.readthedocs.io)[5], and FreeSurfer 6.0[6]. These images were already corrected for motion using FSL's EDDY[7] as part of the HCP minimally-preprocessed diffusion pipeline[8]. We estimated the multi-shell multi-tissue response function using constrained spherical deconvolution[9] and segmented T1-weighted (T1w) images, which were already coregistered to the b0 volume, using the FAST algorithm[10]. We employed anatomically-constrained tractography to generate an initial tractogram with 10 million streamlines using second-order integration over fiber orientation distributions[11]. Subsequently, we applied the spherical-deconvolution informed filtering of tractograms (SIFT2) methodology to yield more biologically accurate measures of fiber connectivity[12].

To define brain regions or network nodes, we utilized the 200-region atlas developed by Schaefer et al. This atlas was mapped to each individual's FreeSurfer surfaces using spherical registration[13]. Additionally, this atlas provided categorical assignments of regions into seven canonical functional brain networks (Visual network: VN, Somatomotor network: SMN, Dorsal attention network: DAN, Anterior salience network: SN, Limbic network: LN, Fronto-parietal network: FPN, Default mode network: DMN).

By combining this atlas with the filtered streamlines, we derived 200 × 200 anatomical connectivity matrices. These matrices contained information on the number of streamlines and fiber length connecting each pair of regions. We then averaged these connectomes across the 400 HCP subjects, resulting in a population-representative connectome matrix.

To prepare this matrix for physiological network modeling, we rescaled the values. This rescaling involved two steps: first, taking the matrix Laplacian, which set each row sum (i.e., each node's weighted in-degree) to zero by subtracting row sums from the diagonal; and second, performing scalar division of all entries by the matrix norm. This ensured the linear stability of the matrix, with all eigenvalues having a negative real part except for one eigenvalue, which was zero. The Laplacian approach has been commonly employed in previous research on whole-brain modeling[14–16].

##### 1.3 Large-scale connectome-based neurophysiological brain network model

As previously outlined, our brain network model encompasses 200 cortical areas, each representing the population-averaged activity of an individual brain region in line with mean-field theory principles[17]. In describing the activity at each node, we employed the Jansen-Rit (JR) equations, a widely adopted neurophysiological model applied to both stimulus-evoked and resting-state EEG activity measurements[18–20], that we have also recently employed for modelling TMS-evoked responses[2].

The JR model serves as a relatively coarse-grained representation of the cortical microcircuit, consisting of three interconnected neural populations: pyramidal projection neurons, excitatory interneurons, and inhibitory interneurons. While the excitatory and inhibitory populations both receive input from, and provide feedback to the pyramidal population, they do not interact with each other directly. Consequently, the circuit motif (Fig. 4C) comprises one positive and one negative feedback loop. Within each of these three neural populations, the post-synaptic somatic and dendritic membrane response to incoming action potentials is described by second-order differential equations.

$$\ddot{v}\left(t\right)+\frac{2}{\tau_{e,i}}\dot{v}\left(t\right)+\frac{1}{\tau_{e,i}^{2}}v\left(t\right)=\frac{H_{e,i}}{\tau_{e,i}}m\left(t\right) \quad (1)$$

which is equivalent to a convolution of incoming activity with a synaptic impulse response function

$$v(t)=\int_{0}^{\infty}d\tau m(\tau)\cdot h_{e,i}(t-\tau) \quad (2)$$

whose kernel $h_{e,i}$ is given by

$$h_{e,i}=\frac{H_{e,i}}{\tau_{e,i}}\cdot t\cdot exp(-\frac{t}{\tau_{e,i}}) \quad (3)$$

where $m$ is the (population-average) presynaptic input, $v$ is the postsynaptic membrane potential, $H_{e,i}$ is the maximum postsynaptic potential, and $\tau_{e,i}$ a lumped representation of delays occurring during the synaptic transmission.

The synaptic response function, often referred to as a pulse-to-wave operator following Freeman's terminology[21], plays a crucial role in regulating the excitability of the neural population. Of specific relevance to our current study are the time constants $\tau_e$ and $\tau_i$, which govern this excitability.

In addition to the pulse-to-wave operator for synaptic responses, each neural population incorporates a wave-to-pulse operator that determines the population's output, represented by the (population-average) instantaneous firing rate. The firing rate is a function of the somatic membrane potential and follows a sigmoidal pattern,

$$S(v) = \frac{e_0}{1 - exp(r(v_0 - v))} \quad (4)$$

where $e_0$ is the maximum firing rate, $r$ is the steepness of the sigmoid function, and $v_0$ is the postsynaptic potential for which half of the maximum firing rate is achieved.

In practice, as is standard with the JR model, we express the three sets of second-order differential equations that follow the structure of Equation X1, as three sets of coupled first-order differential equations. As a result, the complete JR system for each individual cortical area, denoted as j ∈ {i,..., N} within our network comprising N=200 regions, can be represented by the following six equations:

$$\dot{v}_{j1} = x_{j1} \quad (5)$$

$$\dot{x}_{j1} = \frac{H_e}{\tau_e}\left(p_j + \mathrm{conn}_j\right) + C_1 S(C_2 v_{j3}) - \frac{2}{\tau_e} x_{j1} - \frac{1}{\tau_e^2} v_{j1} \quad (6)$$

$$\dot{v}_{j2} = x_{j2} \quad (7)$$

$$\dot{x}_{j2} = \frac{H_i}{\tau_i}\left(C_4 S(C_3 v_{3j})\right) - \frac{2}{\tau_i} x_{j2} - \frac{1}{\tau_i^2} v_{j2} \quad (8)$$

$$\dot{v}_{j3} = x_{j3} \quad (9)$$

$$\dot{x}_{j3} = \frac{H_e}{\tau_e}\left(S(v_{j1} - v_{j2})\right) - \frac{2}{\tau_e} x_{j3} - \frac{1}{\tau_e^2} v_{j3} \quad (10)$$

where $v_{1,2,3}$ is the average postsynaptic membrane potential of the excitatory interneuron, inhibitory interneuron, and pyramidal cell populations, respectively. The input from other nodes in the whole-brain network

$$conn_j(t) = S(\sum_{k \neq j} a_{jk} x_{k1}(t - m_{jk}) \quad (11)$$

In this context, $a_{jk}$ represents the $j_{th}$ row and $k_{th}$ column within the connectivity matrix A, which, in our case, corresponds to the rescaled connectivity Laplacian as previously described. The variable $conn_j$ is exclusively involved in the excitatory population and serves to aggregate excitatory activity from other nodes within the network.

Importantly, due to the finite velocity of long-range axonal conduction, these inputs exhibit delays ranging from approximately 5 to 50 ms, with variations specific to each connection depending on their physical length. These temporal lags are determined by $m_{jk}$, denoting the $j,k_{th}$ entry in the delays matrix $M$, calculated as $T / s$. Here, $T$ represents the inter-regional fiber tract length matrix, and s denotes the global axonal conduction velocity.

Of particular significance in the present work, we modeled the stimulus-evoked depolarization of the resting membrane potential by introducing an external perturbing voltage offset, denoted as $p_j$, which is applied to the excitatory interneuron population.

We used the intracerebral sEEG and threshold-based approach to identify brain regions that were significantly affected by the applied stimulus, giving them non-zero entries in $p_j$. To do this, we calculated the mean and standard deviation of the absolute evoked data for the time window around the stimulus (10ms before and after the stimulus was delivered). A threshold was set at two times the standard deviation above the mean, with brain regions whose sEEG signal exceeded this threshold within that window considered to be regions directly influenced by the stimulus. To quantify the degree of stimulation in each identified region, we implemented a distance-based weighting scheme. We computed a distance matrix that described the spatial relationships between the ROIs. This matrix enabled us to rank the ROIs based on their proximity to the stimulated regions. The ROIs closest to the stimulated areas were assigned higher weights, indicating a greater influence of the stimulus.

The channel-level hd-EEG signals in our model were derived by computing the difference between the excitatory $(v_1(t))$ and inhibitory $(v_2(t))$ interneuron post-synaptic potentials (corresponding to the net somatic membrane potential of the pyramidal cell population[22]) at each cortical parcel, and then projecting these signals into the hd-EEG channel space[23].

$$Y = G \cdot X + \epsilon \quad (12)$$

where $Y$ is the channel-level hd-EEG signals and $G$ is the lead field matrix derived by averaging the sensor-level forward solution over cortical regions.

##### 1.4 Individual-subject Jansen-Rit connectome model parameter estimation from hd-EEG data

We employed a recent technique[2,24] for optimizing parameters in our brain network model when fitting individual-subject evoked potential waveforms and extracting subject-specific physiological parameters from empirical data. A notable component of this methodology is our implementation of the connectome-base neural mass model described above in PyTorch[25], a machine learning software library that is widely used in both academic and commercial sectors. In order to systematize our and others' work with this novel approach, we have recently developed a new Python library, Whole-Brain Modelling in PyTorch (WhoBPyT), available at github.com/griffithslab/whobpyt, which implements and documents the basic technique and our applications of it to resting and stimulus/task-evoked fMRI, EEG, MEG, sEEG and fNIRS data.

Transitioning to the PyTorch-based framework from more conventional numerical simulation libraries required some minor adjustments to accommodate tensor data structures and greater memory load. However, this shift offers a significant advantage by seamlessly accommodating gradient-based parameter optimization through automatic differentiation-based algorithms. This is particularly useful for handling complex sets of equations that do not yield easily computable Jacobians. Our approach falls in line with a growing trend[26,27], where the parallels between physiologically-based large-scale brain network models and deep recurrent neural networks in machine learning prove both technically and conceptually valuable.

Recently, we have successfully applied this technique to fast-timescale evoked responses to dissect the spatio-temporal physiological origin of TMS-evoked potentials[2].

The optimization algorithm operates by segmenting a subject's multi-channel evoked potential waveform, which spans 400 ms (from -100 ms to +300 ms post-stimulus) and is trial-averaged, into discrete, non-overlapping windows of 20 ms, referred to as "batches." The entire simulation covers a duration of 400 ms, encompassing a 100 ms baseline period before the electric pulse and a 300 ms period after the introduction of stimulation. Prior to the commencement of the 100 ms baseline, a 20 ms burn-in period is included to allow the system to stabilize following the initial transient caused by randomly assigned initial conditions for the state variables.

Iterating through each batch in the time series sequentially, we generated the JR model-simulated evoked potentials ŷ using the current parameter values and forward-Euler integration of equations 5-10. We then assessed its alignment with the empirical evoked potentials y by employing the mean-squared error (MSE) loss function

$$L = \frac{1}{N_t} \sum_{t=1}^{N_t} \left( \frac{1}{N_{ch}} \sum_{i=1}^{N_{ch}} \left( \mathbf{y}_i(t) - \hat{\mathbf{y}}_i(t) \right)^2 \right) \quad (13)$$

where $N_t$ is the number of the time points, $N_{ch}$ is the number of hd-EEG channels, and $L$ is the total loss. It is assumed that the model parameters are Gaussian. Together with a complexity-penalizing regularization term on each model parameter $\theta$,

$$C = \ln \sigma + \frac{1}{\sigma^2} (\theta - \mu)^2 \quad (14)$$

Here, the mean *(μ)* and standard deviation *(σ)* of the model parameter $\theta$ serve as optimization hyperparameters. Equation 14 characterizes the complexity of the model parameters, serving as a regularization term to prevent overfitting and enhance model robustness. The loss function $L$ and the complexity term $C$ are combined into a final objective function, which is then supplied to PyTorch's native stochastic gradient descent-based algorithm ADAM[28]. As the batch window traverses the entirety of the evoked potential time series, it cyclically reverts to the beginning and repeats until convergence is achieved. Upon optimization completion, the last 100 batches are used to compute the average value for each parameter, which, in turn, informs subsequent simulations.

We estimated five sets of parameters for each subject: i) physiological parameters defining the dynamic regime of the JR cortical microcircuit, ii) connection weight modulations, iii) lead field modulations, iv) conduction delays, v) hyperparameters. The first of these is the most important as it determines the waveform shape and oscillatory characteristics of the stimulation responses, and in the present context pertains to individual differences in baseline neurophysiology. The second allows for individual variation away from the tractography-based population-representative prior values on anatomical connection strengths described above. It is however important here to strike a balance between empirically-motivated constraints and flexibility during fitting. Our strategy here is to require that the final fitted values (posterior parameter estimates) should remain relatively close to the anatomical connectivity priors, and retain the overall characteristics of that matrix in terms of weight distributions and other spatial features (see discussion of this in [2]). This is achieved by assigning a strong prior variance of 1/50 to each connection weight. Importantly, after fitting the model to each subject's data, we conducted a visual inspection to verify that the posterior mean connection weight matrices preserved the essential topological features observed in empirical neuroimaging data.

Similarly, the projection matrix from sources to channels, computed by solving the Maxwell equations with a boundary element method[29] using MNE[30], begins with a set of population-representative priors, and is allowed to vary per subject with tight constraints by setting small prior variance. For a comprehensive overview of the distribution of the optimized model parameters, please consult Fig. S1.

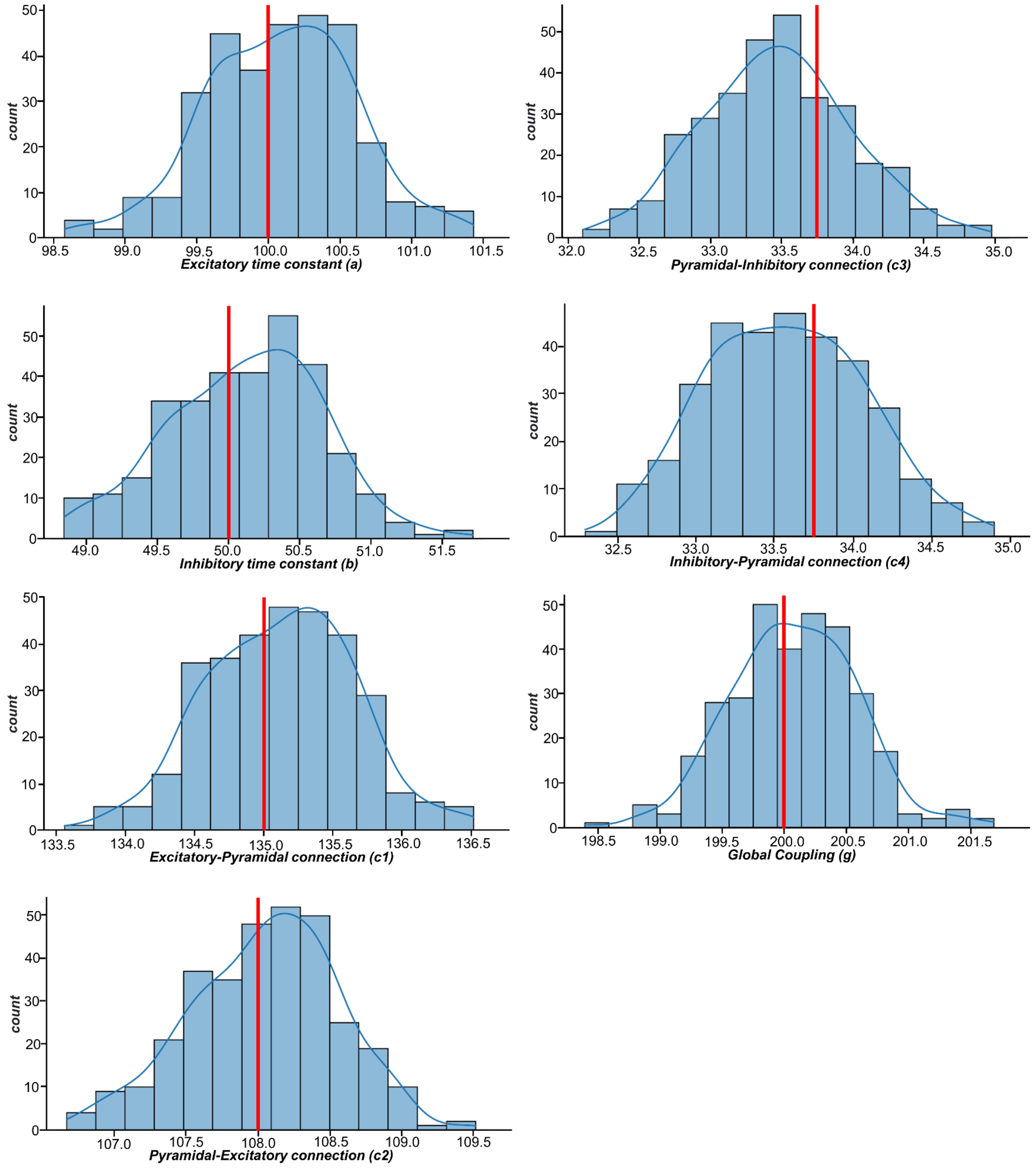


**Fig. S1. Distributions of physiological parameter estimates over subjects.** Histograms and kernel density estimates of the optimized values for the Jansen-Rit model physiological parameters over all subjects. Also shown are prior values for every parameter (vertical red lines). Parameter estimation was performed using our novel automatic differentiation and gradient-based approach inspired by current techniques in deep learning[2,24].

#### 2. SUPPLEMENTARY RESULTS

##### 2.1 Network differences in evoked power are associated with late low frequency

As shown in Fig. S2 Time-frequency analysis on the scalp hd-EEG evoked GFP data revealed two significant clusters (cluster#1: from 3ms to 75ms - average frequency: 16.76Hz; cluster#2: from 65ms to 186ms - average frequency: 5.51Hz). This indicates that the perturbation induces specific changes to spontaneous rhythmic activity at high-alpha (early response) and theta (late response) frequencies. Moreover, when again breaking these results down according to the stimulated network, Wilcoxon-Mann-Whitney U pairwise comparisons showed significant differences in evoked power for both cluster#1: FPN-LN: W=665.5, p=0.01; FPN-VSN: W=455, p=0.005; FPN-SMN: W=1464.5, p<00001; DMN-SMN: W=3565.5, p=0.0001; DMN-VN: W=1130, p=0.01; SN-SMN: W=3483.5, p=0.001; and cluster#2: FPN-VSN: W=428, p=0.02; FPN-SMN: W=1357, p=0002; DMN-SMN: W=3318, p=0.005; DMN-VN: W=1052 , p=0.03). This indicates that the stimulus-evoked power was significantly higher for high-order networks and specifically for late responses (cluster#2).

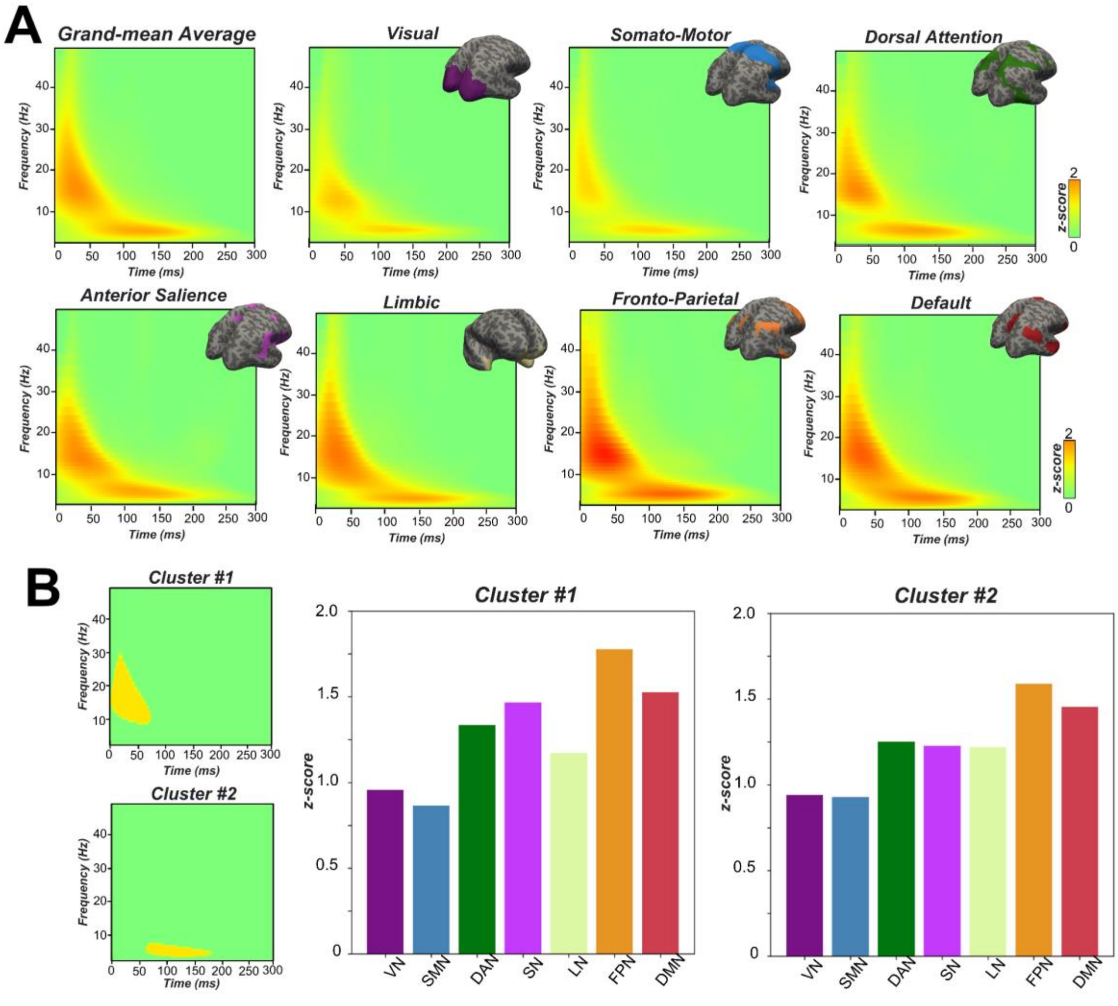


**Fig. S2. Empirical hd-EEG evoked time-frequency spectra (A)** Evoked time-frequency spectral responses as a function of stimulated network. Comparing evoked power over the 300ms window against baseline across all stimulation sites, two distinct significant clusters emerged: Cluster #1 (from 3ms to 75ms, average frequency: 16.76Hz), representing early and transient responses in the high-alpha range; and Cluster #2 (from 65ms to 186ms, average frequency: 5.51Hz), reflecting a later and more sustained theta-frequency response component. These findings indicate that network perturbation induces two specific and distinct modifications in spontaneous rhythms during these time frames. **(B)** Binary representations depict the locations of Cluster #1 (top) and Cluster #2 (bottom), along with the corresponding average power extracted from these clusters across all stimulation sessions and grouped according to the perturbed networks. Significantly higher evoked power was observed in high-order networks than low-order networks, with the magnitude of the response following the same cortical hierarchy pattern as the GMFP analyses.

##### 2.2 Connectome-based neurophysiological modelling accurately reproduces subject-specific evoked dynamics

As an important preliminary result, extensive testing confirmed that our new connectome-based neurophysiological model of stimulus-evoked responses achieves robust and accurate recovery of measured hd-EEG time series at both the group-average and individual-subject levels. Fig. S3A shows empirical and fitted (i.e. simulated, with optimized physiological parameters) stimulus-evoked EEG waveforms along with selected topography maps for three example subjects. It is visually evident in these figures that the model accurately captures several individually-varying features of the stimulus-evoked time series, such as the timing of the early and late components, and the extent to which they are dominated by left/right and temporal/parietal/frontal channels. For the latter, this can be seen by comparing the line colors in the upper and lower rows of corresponding columns in Fig. S3A, and using the channel location references given by the channel color map on the top left of each topoplot. Pearson correlations between empirical and simulated stimulus-evoked time series confirmed that an excellent goodness-of-fit was observed at the individual channel level, with time-wise permutation tests revealing a significant Pearson correlation coefficient for every electrode (R2=86%, $p<0.0001$). As well as the millisecond-by-millisecond comparisons and the timing of key waveform components, we also assessed the accuracy of the model in capturing holistic time series properties for both early and late responses.

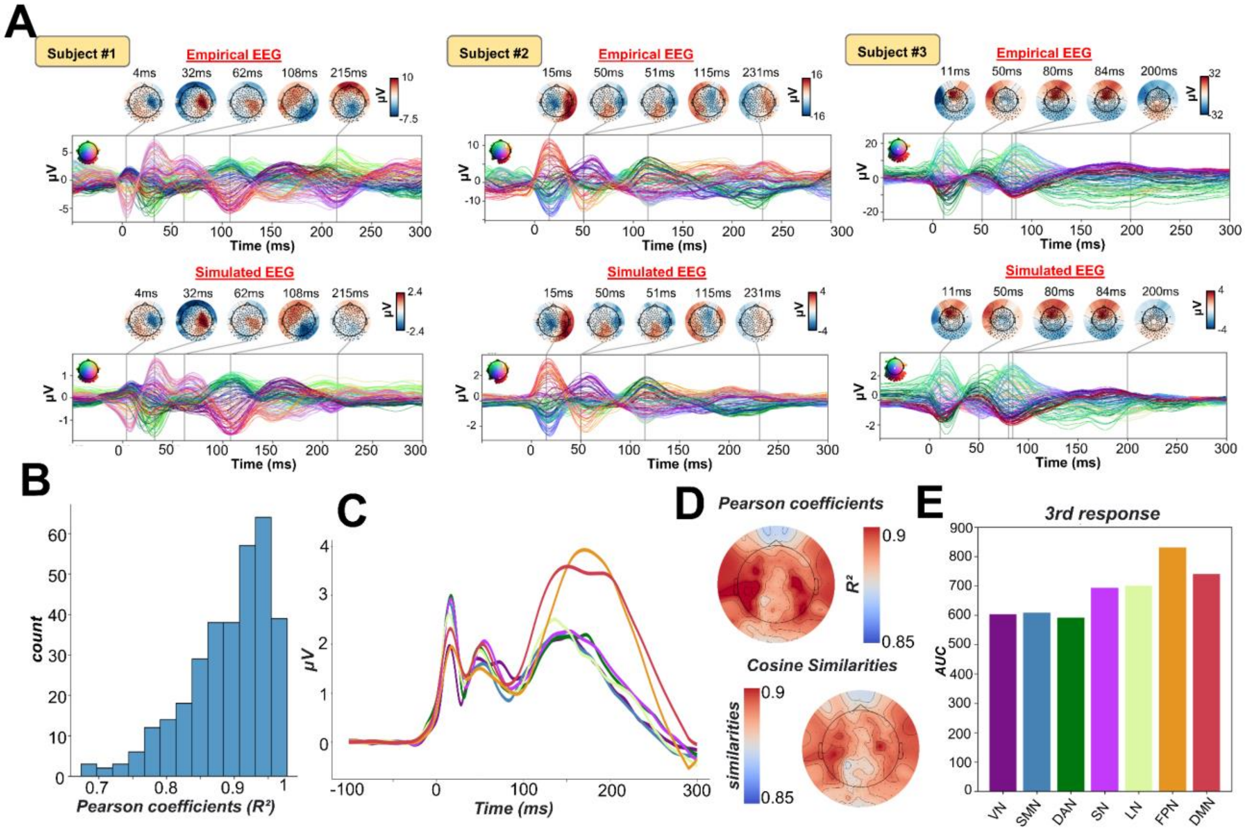


**Fig. S3. Goodness-of-fit between empirical and simulated stimulus-evoked EEG data. (A)** Empirical (upper row) and simulated (lower row) stimulus-evoked EEG butterfly plots, with scalp topographies for three representative subjects. These simulations , showing a robust recovery of individual empirical evoked potential patterns in model-generated activity electroencephalography (EEG) time series. **(B)** Histogram plot showing the distributions of the Pearson correlation coefficients between simulated and empirical time series for each subject and session. **(C)** Simulated GMFP for every stimulated network showing a similar propagation pattern to that shown in Fig. 2 for empirical data. **(D)** Topomaps showing the Pearson correlation coefficients (top - R2=86%, p<0.0001) and cosine similarity (R2=87%, p<0.0001) for every channel,. demonstrating high correspondence between empirical and simulated data. **(E)** Reproduction of the above-described network hierarchy for model-generated data, showing greater activation for high-order regions compared to regions serving primary sensory/motor functions.

#### 3. SUPPLEMENTARY DISCUSSIONS

We have shown that in the time-frequency power spectrum of both the early fast (alpha) and the late slow (theta) evoked waves, high-order networks have a stronger evoked activity, particularly evident for later theta-frequency responses. This is consistent with the view that the individual anatomical structure of a brain area shapes its rhythmic neuronal activity, with the exact dominant frequency of oscillations changing systematically and globally over hierarchical spatial gradients[31]. Indeed, previous MEG resting-state studies found that the dominant peak frequency in a brain area decreases significantly, gradually, and robustly along the posterior-anterior axis, following the global cortical hierarchy from early sensory to higher order areas[32–34]. These findings have established a frequency gradient of resting-state brain rhythms that complements previous anatomical studies reporting a posterior-anterior gradient in microscale and macroscale anatomical features of animal and human cortex[35]. Moreover, stimulus-evoked studies have consistently demonstrated that each cortical area tends to preserve its own natural frequency, even when indirectly engaged by external perturbations through brain connections or stimulated at different intensities[36,37]. Additional supporting evidence from intracranial studies have shown that oscillations typically propagate in a direction from the posterior to the anterior. This directional pattern is attributed to the coordination driven by an overall reduction in intrinsic oscillation frequency from posterior to anterior regions, suggesting a gradual decline in power-frequency along the sagittal axis[38]. This decrease in the posterior-anterior direction has been related to a representational gradient of spatial scales from coarse to fine[39]. In line with this evidence, we have demonstrated here using stimulation-evoked-electrophysiological data that the evoked-oscillations are slower in higher-order regions. This highlights first how regions' evoked responses depend on their position along the cortical gradient, and second - in line with converging evidence across recording methods, species and cortical domain[39] - that representations become more integrated with decreasing dominant frequency of the underlying neuronal population. Interestingly, many studies have demonstrated that faster oscillations underlie feedforward processing, while slower oscillations serve as relays of feedback processing[40–43]. In this framework, the stronger slow wave oscillations in the high-order networks for the late responses might reflect a slower propagation pattern of the feedback wave, which may in turn contribute to slower integration of inputs across multiple recent sensory stimuli.

#### 4. SUPPLEMENTARY REFERENCES

1. Pereira, A. C. *et al.* An in vivo correlate of exercise-induced neurogenesis in the adult dentate gyrus. *Proc. Natl. Acad. Sci. U. S. A.* **104**, 5638–5643 (2007).
2. Momi, D., Wang, Z. & Griffiths, J. D. TMS-evoked responses are driven by recurrent large-scale network dynamics. *eLife* **12**, e83232 (2023).
3. Van Essen, D. C. *et al.* The Human Connectome Project: a data acquisition perspective. *NeuroImage* **62**, 2222–2231 (2012).
4. Jenkinson, M., Beckmann, C. F., Behrens, T. E. J., Woolrich, M. W. & Smith, S. M. FSL. *NeuroImage* **62**, 782–790 (2012).
5. Tournier, J.-D., Calamante, F. & Connelly, A. MRtrix: Diffusion tractography in crossing fiber regions. *Int. J. Imaging Syst. Technol.* **22**, 53–66 (2012).
6. Fischl, B. FreeSurfer. *NeuroImage* **62**, 774–781 (2012).
7. Andersson, J. L. R. & Sotiropoulos, S. N. An integrated approach to correction for off-resonance effects and subject movement in diffusion MR imaging. *NeuroImage* **125**, 1063–1078 (2016).
8. Glasser, M. F. *et al.* The minimal preprocessing pipelines for the Human Connectome Project. *NeuroImage* **80**, 105–124 (2013).
9. Christiaens, D. *et al.* Global tractography of multi-shell diffusion-weighted imaging data using a multi-tissue model. *NeuroImage* **123**, 89–101 (2015).
10. Zhang, Y., Brady, M. & Smith, S. Segmentation of brain MR images through a hidden Markov random field model and the expectation-maximization algorithm. *IEEE Trans. Med. Imaging* **20**, 45–57 (2001).
11. Tournier, J.-D., Calamante, F. & Connelly, A. Improved probabilistic streamlines tractography by 2nd order integration over fibre orientation distributions. *Proc Intl Soc Mag Reson Med ISMRM* **18**, (2010).
12. Smith, R. E., Tournier, J.-D., Calamante, F. & Connelly, A. SIFT2: Enabling dense quantitative assessment of brain white matter connectivity using streamlines tractography. *NeuroImage* **119**, 338–351 (2015).
13. Fischl, B., Sereno, M. I. & Dale, A. M. Cortical surface-based analysis. II: Inflation, flattening, and a surface-based coordinate system. *NeuroImage* **9**, 195–207 (1999).
14. Abdelnour, F., Dayan, M., Devinsky, O., Thesen, T. & Raj, A. Functional brain connectivity is predictable from anatomic network's Laplacian eigen-structure. *NeuroImage* **172**, 728–739 (2018).
15. Atasoy, S., Donnelly, I. & Pearson, J. Human brain networks function in connectome-specific harmonic waves. *Nat. Commun.* **7**, 10340 (2016).
16. Raj, A., Verma, P. & Nagarajan, S. Structure-function models of temporal, spatial, and spectral characteristics of non-invasive whole brain functional imaging. (2022) doi:10.3389/fnins.2022.959557.
17. Deco, G., Jirsa, V. K., Robinson, P. A., Breakspear, M. & Friston, K. The Dynamic Brain: From Spiking Neurons to Neural Masses and Cortical Fields. *PLOS Comput. Biol.* **4**, e1000092 (2008).
18. David, O., Harrison, L. & Friston, K. J. Modelling event-related responses in the brain. *NeuroImage* **25**, 756–770 (2005).
19. Jansen, B. H. & Rit, V. G. Electroencephalogram and visual evoked potential generation in a mathematical model of coupled cortical columns. *Biol. Cybern.* **73**, 357–366 (1995).
20. Spiegler, A., Kiebel, S. J., Atay, F. M. & Knösche, T. R. Bifurcation analysis of neural mass models: Impact of extrinsic inputs and dendritic time constants. *NeuroImage* **52**, 1041–1058 (2010).
21. Freeman, W. J. Mass Action in the nervous system. (1975).
22. David, O., Kilner, J. M. & Friston, K. J. Mechanisms of evoked and induced responses in MEG/EEG. *NeuroImage* **31**, 1580–1591 (2006).
23. Endo, H., Hiroe, N. & Yamashita, O. Evaluation of Resting Spatio-Temporal Dynamics of a Neural Mass Model Using Resting fMRI Connectivity and EEG Microstates. *Front. Comput. Neurosci.* **13**, (2020).
24. Griffiths, J. D. *et al.* Deep Learning-Based Parameter Estimation for Neurophysiological Models of Neuroimaging Data. 2022.05.19.492664 Preprint at https://doi.org/10.1101/2022.05.19.492664 (2022).
25. Paszke, A. *et al. PyTorch: An Imperative Style, High-Performance Deep Learning Library*. http://arxiv.org/abs/1912.01703 (2019) doi:10.48550/arXiv.1912.01703.
26. Richards, B. A. *et al.* A deep learning framework for neuroscience. *Nat. Neurosci.* **22**, 1761–1770 (2019).
27. Suárez, L. E., Richards, B. A., Lajoie, G. & Misic, B. Learning function from structure in neuromorphic networks. 2020.11.10.350876 Preprint at https://doi.org/10.1101/2020.11.10.350876 (2020).
28. Kingma, D. P. & Ba, J. Adam: A Method for Stochastic Optimization. *ArXiv14126980 Cs* (2017).
29. Cuffin, B. N. A method for localizing EEG sources in realistic head models. *IEEE Trans. Biomed. Eng.* **42**, 68–71 (1995).

30. Gramfort, A. *et al.* MNE software for processing MEG and EEG data. *NeuroImage* **86**, 446–460 (2014).
31. Mahjoory, K., Schoffelen, J.-M., Keitel, A. & Gross, J. The frequency gradient of human resting-state brain oscillations follows cortical hierarchies. *eLife* **9**, e53715 (2020).
32. Mellem, M. S., Wohltjen, S., Gotts, S. J., Ghuman, A. S. & Martin, A. Intrinsic frequency biases and profiles across human cortex. *J. Neurophysiol.* **118**, 2853–2864 (2017).
33. Keitel, A. & Gross, J. Individual Human Brain Areas Can Be Identified from Their Characteristic Spectral Activation Fingerprints. *PLoS Biol.* **14**, e1002498 (2016).
34. Hillebrand, A. *et al.* Direction of information flow in large-scale resting-state networks is frequency-dependent. *Proc. Natl. Acad. Sci.* **113**, 3867–3872 (2016).
35. Huntenburg, J. M., Bazin, P.-L. & Margulies, D. S. Large-Scale Gradients in Human Cortical Organization. *Trends Cogn. Sci.* **22**, 21–31 (2018).
36. Rosanova, M. *et al.* Natural Frequencies of Human Corticothalamic Circuits. *J. Neurosci.* **29**, 7679–7685 (2009).
37. Bai, Y., Xuan, J., Jia, S. & Ziemann, U. TMS of parietal and occipital cortex locked to spontaneous transient large-scale brain states enhances natural oscillations in EEG. *Brain Stimulat.* **16**, 1588–1597 (2023).
38. Zhang, H., Watrous, A. J., Patel, A. & Jacobs, J. Theta and Alpha Oscillations Are Traveling Waves in the Human Neocortex. *Neuron* **98**, 1269-1281.e4 (2018).
39. Muller, L., Chavane, F., Reynolds, J. & Sejnowski, T. J. Cortical travelling waves: mechanisms and computational principles. *Nat. Rev. Neurosci.* **19**, 255–268 (2018).
40. Bastos, A. M. *et al.* Visual areas exert feedforward and feedback influences through distinct frequency channels. *Neuron* **85**, 390–401 (2015).
41. van Ede, F., van Pelt, S., Fries, P. & Maris, E. Both ongoing alpha and visually induced gamma oscillations show reliable diversity in their across-site phase-relations. *J. Neurophysiol.* **113**, 1556–1563 (2015).
42. Michalareas, G. *et al.* Alpha-Beta and Gamma Rhythms Subserve Feedback and Feedforward Influences among Human Visual Cortical Areas. *Neuron* **89**, 384–397 (2016).
43. Aggarwal, A. *et al.* Visual evoked feedforward–feedback traveling waves organize neural activity across the cortical hierarchy in mice. *Nat. Commun.* **13**, 4754 (2022).
